# Bacterial and metabolic factors of staphylococcal planktonic and biofilm environments differentially regulate macrophage immune activation

**DOI:** 10.1101/2021.07.26.453923

**Authors:** Elisabeth Seebach, Tabea Elschner, Franziska V. Kraus, Margarida Souto-Carneiro, Katharina F. Kubatzky

**Author notes:** **Correspondence:** Elisabeth Seebach, Katharina F. Kubatzky.

## Abstract

Biofilm formation is a leading cause for chronic implant-related bone infections as biofilms shield bacteria against the immune system and antibiotics. Additionally, biofilms generate a metabolic microenvironment that shifts the immune response towards tolerance. Here, we compared the impact of the metabolite profile of bacterial environments on macrophage immune activation using *Staphylococcus aureus* (SA) and *epidermidis* (SE) conditioned media (CM) of planktonic and biofilm cultures. The biofilm environment had reduced glucose and increased lactate concentrations. Moreover, the expression of typical immune activation markers on macrophages was reduced in the biofilm environment compared to the respective planktonic CM. However, all CM caused a predominantly pro-inflammatory macrophage cytokine response with a comparable induction of *Tnfa* expression. In biofilm CM this was accompanied by higher levels of anti-inflammatory *Il10*. Planktonic CM on the other hand, induced an IRF-7 mediated *Ifnb* expression which was absent in the biofilm environments. For SA but not for SE planktonic CM, this was accompanied by IRF3 activation. Stimulation of macrophages with TLR-2/-9 ligands under varying metabolic conditions revealed that, like in the biofilm setting, low glucose concentration reduced the *Tnfa* to *Il10* mRNA ratio. However, addition of extracellular L-lactate but not D-lactate increased the *Tnfa* to *Il10* mRNA ratio upon TLR-2/-9 stimulation. In summary, our data indicate that the mechanisms behind the activation of macrophages differs between planktonic and biofilm environments. These differences are independent of the metabolite profiles, suggesting that the production of different bacterial factors is ultimately more important than the concentrations of glucose and lactate in the environment.

## Introduction

Chronic implant-related bone infections are a major complication in orthopedic and trauma surgery with severe consequences for the patients including long-term antibiotic treatment, repeated surgeries, implant revision, and at worst amputation of the infected limb [1, 2]. Bacteria can take advantage of the indwelled, foreign body and form biofilms on the implant surface. Such infections are frequently persistent because the biofilm matrix acts as a physical barrier that shields bacteria against eradication through host immune system and antibiotic treatment [3–6]. Furthermore, several studies show that biofilm formation promotes a more tolerant immune response, which facilitates bacterial persistence [7, 8].

*Staphylococcus aureus* (SA) and *epidermidis* (SE) are the most frequently isolated bacteria in implant-related bone infections [1, 6]. SA expresses a broad range of virulence factors, is able to form small colony variants (SCV), can survive inside osteoblasts and osteoclasts [9, 10], persist in cortical bone structures [11], and form biofilms [12]. The commensal SE in comparison mainly relies on biofilm formation as an immune evasion strategy but does not produce aggressive pathogenic factors [13, 14]. Thus, SA is pre-dominantly found in early and acute infections that are associated with pain, swelling, and fever and implicate a high risk for infection recurrence after antibiotic and surgical treatment [15]. SE causes only mild symptoms and low-grade inflammation. Therefore, SE infections are commonly detected when the infection gets chronic, which is closely linked to biofilm formation [16].

Recognition of specific bacterial pathogen associated molecular patterns (PAMPs) by innate immune cells causes activation of pattern recognition receptors (PRRs), such as the toll-like receptors (TLRs). Binding of a PAMP to its respective TLR leads to the activation of the nuclear factor ‘kappa-light-chain-enhancer’ of activated B cells (NF-κB) pathway and subsequent to the induction of inflammatory cytokines. Further, TLR activation can induce a Type 1 IFN response mainly via the IRF7 pathway resulting in the production of IFN-α and IFN-β [17]. Next to TLRs, cytosolic PRRs including Nod-like receptors (NLRs) and dsDNA sensors such as cGAS/STING contribute to an effective immune response against invading bacteria [18, 19]. Binding of SA lipopeptides and lipoteichoic acid (LTA) triggers immune cell activation through surface-bound TLR-2 while the recognition of bacterial DNA motifs occurs via endosomal TLR-9 [20]. Biofilm formation however, is discussed to prevent TLR-2 and TLR-9 recognition of embedded bacteria by masking or retaining PAMPs within the biofilm matrix [21, 22].

Macrophages are innate immune cells with an important role in the first line of defense against invading pathogens. Together with the other cells of the innate immune system, they fight bacterial infections by phagocytosis and production of anti-microbial molecules such as nitric oxygen (NO), anti-microbial peptides, and cytokines [23]. *In vitro*, macrophages can be polarized into rather pro-inflammatory (M1) or anti-inflammatory (M2) phenotypes [24], associated either with bacterial clearance or tolerance and persistence, respectively [25]. This classification is defined by the expression of specific surface markers, inducible nitric oxide synthase (iNOS, M1) or arginase 1 (Arg-1, M2), and the cytokine profile of the respective macrophage population (M1: TNF-α, IL-1 and IL-12 vs. M2: IL-10 and TGF-β) [25]. Macrophage subtypes are also characterized by different metabolic activities. While M1 macrophages pre-dominantly rely on aerobic glycolysis, M2 macrophages are associated with oxidative phosphorylation (OxPhos) [26]. Usually, an effective pro-inflammatory macrophage immune response is able to clear an infection caused by planktonic bacteria. However, this often fails once these bacteria start building a biofilm. Metabolically, biofilm formation is characterized by extraction of glucose from the environment and accumulation of fermentation products such as lactate due to the anaerobic growth conditions within the biofilm [27]. This metabolic microenvironment is considered to support the biofilm-mediated anti-inflammatory macrophage polarization and thus, impairs them to exert their clearance functions [28].

So far, most studies focused on investigation of the infection after a biofilm has formed. Here, we addressed the question if the differences in metabolite levels could be a central cause for the distinct macrophage response against planktonic and biofilm environments, or if bacterially derived factors impact macrophage activation. Thus, we treated murine RAW 264.7 macrophages with conditioned media (CM) generated from SA or SE planktonic and biofilm cultures, respectively and analyzed the induction of pro- and anti-inflammatory cytokine production and their metabolic activity. Additionally, we evaluated the effect of low glucose or high lactate concentrations on macrophage polarization upon combined TLR-2/-9 activation or stimulation with SA planktonic CM.

## Materials and Methods

### Bacteria culture and preparation of conditioned media

*Staphylococcus aureus* strain ATCC 49230 (UAMS-1, isolated from a patient with chronic osteomyelitis) [29] and *Staphylococcus epidermidis* strain DSM 28319 (RP62A, isolated from a catheter sepsis) were used for preparation of conditioned media. Bacteria were cultured on Columbia agar plates with 5% sheep blood (BD, Germany) and streaked onto fresh agar plates a day before experiment. 3 to 5 colonies were transferred into trypticase soy bouillon (TSB; BD, Germany) and cultivated under shaking for 3 hours at 37 °C to receive growth state bacteria. Bacterial density was measured photometrically (Den-1, Grant Instruments, UK) and adjusted to a concentration of 6*10^5^ CFU/ml in DMEM high glucose (Anprotec, Germany) + 10% heat-inactivated fetal calf serum (FCS; Biochrom GmbH, Germany). For planktonic culture, bacteria were cultivated under shaking (200 rpm) for 24 hours at 37°C and 5% CO2. For biofilm culture, bacteria were plated in 24 well with 1 ml per well and cultivated under static conditions for 6 days. In biofilm cultures, medium carefully was replaced every 24 hours. For CM, planktonic medium after 24 hours of culture or the last 24 hours medium change before day 6 biofilm culture was harvested by centrifugation at 4000 rpm for 15 min at 4°C. For biofilm CM, media of wells were pooled before centrifugation. Harvested media were streaked onto agar plates, cultivated over night at 37°C and bacterial appearance (colony size and color) was controlled to ensure no contamination by other bacteria. Supernatants were then sterile filtered through a 0.2 μm filter and frozen at −80°C. To rule out remaining bacterial growth, sterile filtered media were inoculated in TSB and cultivated over night at 37°C. Before use in cell culture, pH of CM was adjusted to physiological pH of growth medium (DMEM high glucose + 10% FCS) by drop by drop titration with 0.5 N NaOH and color check of the pH indicator Phenol Red. Aliquots were stored at −80°C. Planktonic and biofilm CM from the same approach were compared within one experiment. For unstimulated CM control, growth medium (DMEM high glucose + 10% FCS) of the respective approach was treated analogously to CM without bacteria inoculation.

### ^1^H NMR metabolomics

^1^H NMR spectra of one representative CM approach were acquired using a 400 MHz Bruker spectrometer (Bruker Ultrashield™ Plus 400) equipped with a 5-mm indirect detection probe. Each spectrum covered a spectral width of 6.4 kHz. A NOESY1D sequence with water-signal suppression and a 30-degree pulse and a total repetition time of 6.5 seconds were applied to ensure full relaxation of all proton nuclei in the samples. Before Fourier transformation, each free induction decay (FID) was multiplied by a decaying exponential with a decay constant of 0.3 Hz. To allow comparison between different spectra, sodium fumarate (10 mM), dissolved in a 0.2 M phosphate buffer solution prepared with D2O (99.9%), was used as an internal standard. NMR samples consisted of 140 μL of CM plus 35 μL of fumarate standard. Before evaluation, spectra were phase adjusted and baseline corrected using the Bruker TopSpin 3.6.3 software. Spectra were further calibrated by setting the resonance of fumarate to δ = 6.5 parts per million (ppm). For comparison of different CM, spectra were adjusted to each other by equalizing the resonance of fumarate.

### Cell culture and stimulation of macrophages

The murine macrophage cell line RAW 264.7 (ATCC TIB-71, USA) was used for the experiments [30]. RAW 264.7 cells were cultivated in DMEM high glucose + 10% heat-inactivated FCS + 1% Pen/Strep at 37°C and 5% CO_2_. Cells were plated into suitable well plate formats, treated with CM 1:1 diluted in fresh cell growth media or PC (positive control: 1 μg/ml TLR-2 ligand Pam3CSK4 and 100 nM TLR-9 ligand CpG ODN 1668, both InvivoGen, USA). For experiments with different glucose concentrations, cells were cultivated on passage before experiment and stimulation was done either in high glucose DMEM (4.5 g/l) or low glucose DMEM (1 g/l, Anprotec, Germany). In experiments with different extracellular lactate concentration, sodium L- or D-lactate (10, 15 and 20 mM, both Sigma-Aldrich, Germany) were added simultaneously to stimulation.

### Flow cytometry

For FACS analysis, 2 million cells / well were plated with 1 ml fresh growth media and 1 ml CM. After 20 hours supernatants were frozen at −80°C for further investigation and cells were washed twice with cold PBS. For surface marker staining, 100 μl of cell suspension was either left in PBS / 2% BSA for the unstained control or stained with 0.2 mg/ml FITC anti-TLR-2 (Novus Biologicals, UK), PE anti-MHC II (Invitrogen, USA) or PE anti-CD80 (BioLegend, USA) antibodies at 4°C for 1 hour. Cells were washed two times in cold PBS, resuspended in 150 μl cold PBS and then analyzed with the BD FACSCanto™ Flow Cytometer (BD Biosciences, USA). After measurement, unstained controls were additionally incubated with 30 nM Sytox^R^ Green nucleic acid stain (Invitrogen, USA) for 5 min and live / dead contribution was recorded. For intracellular staining of TLR-9, 500 μl of cell suspension were combined with 500 μl fixation buffer (BioLegend, USA), incubated at 37°C for 15 min, centrifuged at 2000 rpm for 5 min, 2x washed with PBS / 5% BSA and then stored at 4°C, over-night in PBS / 5% BSA. Next day, cells were centrifuged and resuspended in 1 ml of −20°C cold TruePhos™ Perm Buffer (BioLegend, USA) and incubated at −20°C for 1 hour. Cells were then washed two times and resuspended in 500 μl PBS / 2% BSA. 100 μl of cell suspension was either kept unstained, stained with 0.5 mg/ml Alexa647 anti-TLR-9 antibody (Novus Biologicals, UK) or respective isotype at RT for 30 min. Cells were washed twice, resuspended in 300 μl PBS / 2% BSA and analyzed with the BD FACSCanto™ Flow Cytometer. Results were further analysed using the Flowing Software (version 2.5.1, Turku Bioscience, Finland).

### Cytometric bead array

Supernatants of FACS surface marker analysis were used for cytometric bead array (CBA, LegendPlex, BioLegend, USA) according to the manufacturer’s protocol. A Mouse Inflammation Panel (Mix and Match Subpanel) was used including beads against TNF-α, IL-10 and IFN-β. In short, supernatants were centrifuged, diluted 1:5 with Assay Buffer, standard samples were prepared and samples were transferred into a V-bottom plate. Bead mix was prepared, added to the samples and incubated on a shaker over night at 4°C in the dark. Next day, plate was washed 2 times and incubated with detection antibody for 1 hour at RT while shaking. SA-PE was added and further incubated for 30 min at RT while shaking. Plate was washed two times; bead pellet was resuspended in wash buffer and data acquisition was done with the BD^®^ LSR II Flow Cytometer. Analysis and calculation of cytokine concentrations were performed with the included LEGENDplex™ Data Analysis Software (version 8.0).

### Gene expression analysis

For gene expression analysis, cells were stimulated as indicated in the figure legends. Total RNA extraction was performed using the innuPREP RNA Mini Kit 2.0 (Analytik Jena, Germany) according to the manufacture’s protocol. In short, cells were scraped in lysis buffer and transferred to a DNA elution column. RNA in the lysate was precipitated by adding 70% ethanol, transferred to an RNA column, washed and eluated in H2O. Total RNA concentration was measured using the NanoDrop^®^ ND-1000 spectrophotometer (Thermo Scientific, Germany). 1 μg of total RNA was subjected to cDNA synthesis using the Biozym cDNA synthesis Kit (Biozym Scientific GmbH, Germany) according to the manufacturer’s protocol using Oligo (dT) primer. A noRT sample (w/o Reverse Transcriptase) consisting of pooled total RNA of all samples of one experiment was prepared. cDNA was diluted 1:1 in H_2_O and stored at −20°C. 2 μl cDNA template and 400 nM of respective primer pairs (Table 1) were used in qPCR. mRNA levels were evaluated in a two-step PCR reaction (StepOnePlus Real-Time PCR Cycler, Applied Biosystems, USA) with 60°C annealing/extension temperature for 40 cycles using the 2x qPCRBIO SyGreen Mix Hi-ROX (PCR Biosystems Ltd., UK). Quality of qPCR runs and specificity of qPCR products were controlled by included noRT and water samples for each experiment and primer pair and melting curve comparison. mRNA levels of the respective genes of interest (Table 1) were normalized to the reference gene HPRT-1 and calculated by the 2^-ΔCT^ method.

**Table 1.**
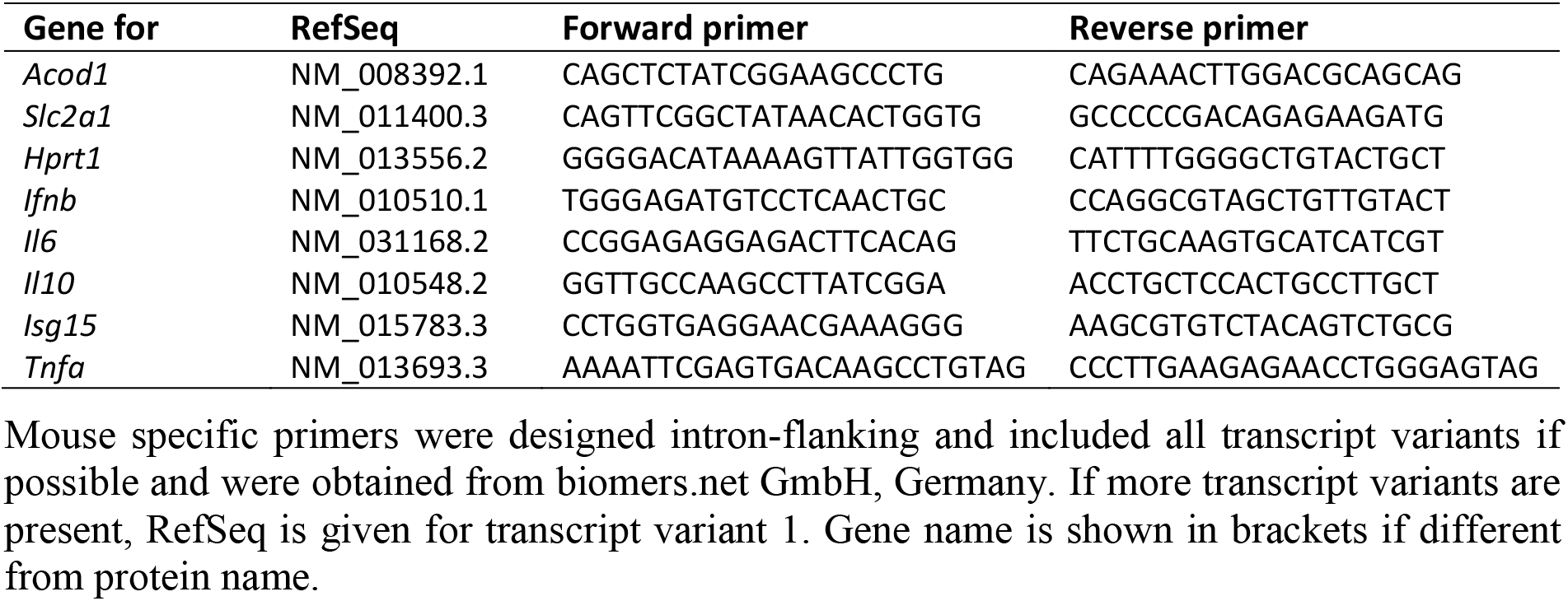
List of oligonucleotides used for quantitative RT-PCR analysis.

### Immunoblotting

For protein analysis by Western Blot, cells were stimulated as indicated in the figure legends. Cells were lysed in RIPA buffer (1% v/v NP-40 (IGEPAL^®^ CA-630), 0.25% sodium deoxycholate, 50 mM Tris pH 8.0, 150 mM NaCl, 1 mM EDTA pH 8.0, 1 mM Na_3_VO_4_, 1 mM NaF) with EDTA-free protease inhibitors (cOmplete™ Tablets) and phosphatase inhibitors (PhosSTOP™, both Roche Diagnostics GmbH, Germany) for 1 hour at 4°C under rotation. Lysates were centrifuged at 14000 rpm for 20 min at 4°C. Protein concentrations were determined by BCA assay (Cyanagen Srl, Italy), samples were adjusted to 10 μg protein per 20 μl with ddH2O and 5 μl 4x SDS sample buffer with 10% β-mercaptoethanol and loaded on pre-cast gradient 4-20% Tris-glycine gels (anamed Elektrophorese GmbH, Germany). Proteins were transferred onto an Amersham™ Protran™ 0.45 μm nitrocellulose membrane (GE Healthcare, UK). Membranes were blocked with BlueBlock PF (Serva Electrophoresis GmbH, Germany) for 30 min at RT before incubation with primary antibodies (Table 2) over night at 4°C. After three times washing with TBST, membranes were incubated with an anti-rabbit HRP-linked secondary antibody (1:1000, Cell Signaling Technology, USA) for 1 hour at RT. Blots were developed with ECL substrate (WESTAR ETA C ULTRA 2.0, Cyanagen Srl, Italy) and imaged in the ChemoStar ECL & Fluorescence Imager (Intas Science Imaging Instruments GmbH, Germany).

**Table 2.** List of antibodies used for Immunoblotting (Western Blot).

| Protein | Source | Size (kDa) | Dilution | Company |
| --- | --- | --- | --- | --- |
| HSP-90 | rabbit | 90 | 1:1000 | Cell Signaling Technology, USA |
| Phospho-IRF3 (Ser396) | rabbit | 45-55 | 1:1000 | Cell Signaling Technology, USA |
| Phospho-IRF7 (Ser437/438) | rabbit | 55 | 1:1000 | Cell Signaling Technology, USA |
| Phospho-NFκB p65 (Ser536) | rabbit | 65 | 1:1000 | Cell Signaling Technology, USA |

Antibodies were all recommended for use in mouse and applied according to the manufacturer's advice. Proteins were detected by chemiluminescent luminol reaction after incubation with respective HRP-linked secondary antibody and imaged in a ChemoStar ECL Imager.

### L-lactate detection

50.000 cells / well were plated in 96 well format and stimulated with 100 μl CM and 100 μl fresh growth media for 24 hours. Supernatants of three replicates were pooled and stored at - 80°C until further processing. L-lactate concentration was measured in CM as well as in supernatants of CM stimulated macrophages using an enzyme-based bioluminescent assay according to the manufacturer’s protocol (Lactate-Glo™ Assay, Promega GmbH, Germany). CM and supernatants were used 1:50 (planktonic CM) or 1:100 (other) diluted with PBS, a standard curve with defined L-lactate concentrations (0 – 200 μM) was included. Samples and standard were incubated with the lactate detection reagent for 1 hour at RT and light emission was recorded by luminometer (LUMIstar^®^ Optima, BMG LABTECH, Germany).

### ATP detection

50.000 cells / well were plated in 96 well format and stimulated with 100 μl CM and 100 μl fresh growth media for 24 hours. Samples were performed in triplicates. Media were removed and 100 μl of CTG reagent (CellTiter-Glo^®^, Promega GmbH, Germany; 1:1 with PBS) per well was added. Cells were lysed for 1 min at RT under continuous shaking. After 10 min incubation in the dark, supernatants were transferred into a white 96 well plate. Relative ATP content was determined by bioluminescent light reaction in a luminometer (LUMIstar^®^ Optima, BMG LABTECH, Germany).

### Mitochondrial activity

300.000 cells / well were transferred in 24 well plates and stimulated with CM 1:1 diluted with fresh growth media in a total of 1 ml. Mitochondrial activity was measured after 24 hours by adding 100 nM of a mitochondrial membrane potential-sensitive dye (stock conc.: 1 mM in DMSO, MitoTracker^®^ Deep Red FM, Cell Signaling Technology, USA) to the cells for 30 min at 37°C and 5% CO2. Cells were then washed three times with cold PBS, scraped in PBS and transferred into FACS tubes. Mitochondrial activity was analyzed with the BD FACSCanto™ Flow Cytometer according to the fluorescence intensity of the dye. Only the living cell population was included in further analysis using the Flowing Software (version 2.5.1, Turku Bioscience, Finland).

### Statistical analysis

Experiments were done in n = 4-5 independent replicates as stated in the figure legends. Data are presented as mean + SD and single values as dots. Statistical evaluation was performed using Ordinary one-way ANOVA with post-hoc multiple comparison testing and Bonferroni correction. A p-value below 0.05 was considered as statistically significant. * is indicating significance against Medium, # is showing significance between different treatments. Data analysis was performed with GraphPad Prism for Windows (Version 9.3.1, GraphPad Software Inc., USA).

## Results

### In vitro generated CM represent the in vivo biofilm environment characterized by low glucose and high lactate levels

To validate the experimental system, we monitored the ability of the two bacterial strains, SA and SE, to form biofilms. As described in the literature, we could reproduce that SA strain UAMS-1 is a moderate biofilm producer on non-coated plastic surfaces *in vitro* [31], whereas SE RP62A is known to possess strong *in vitro* biofilm formation capacities on plastic [32] (Fig. 1A, B). Supernatants were prepared from bacterial cultures under planktonic or biofilm-specific culture conditions (Figure 1B, C). ^1^H-NMR analysis of these supernatants revealed that glucose could still be detected in the CM of planktonic SA and SE, whereas all glucose was metabolized in the respective biofilm CM (Fig. 1D). In all conditions, bacteria released acetate. Interestingly, the levels differed between SA and SE and were higher in CM from planktonic SA and biofilm SE. As lactate was strongly increased in the CM of SA and SE biofilm cultures, but could only be detected in residual amounts in the respective planktonic CM we conclude that the generated CM represent the *in vivo* situation, characterized by glucose deprivation and lactate accumulation.

**Figure 1.**
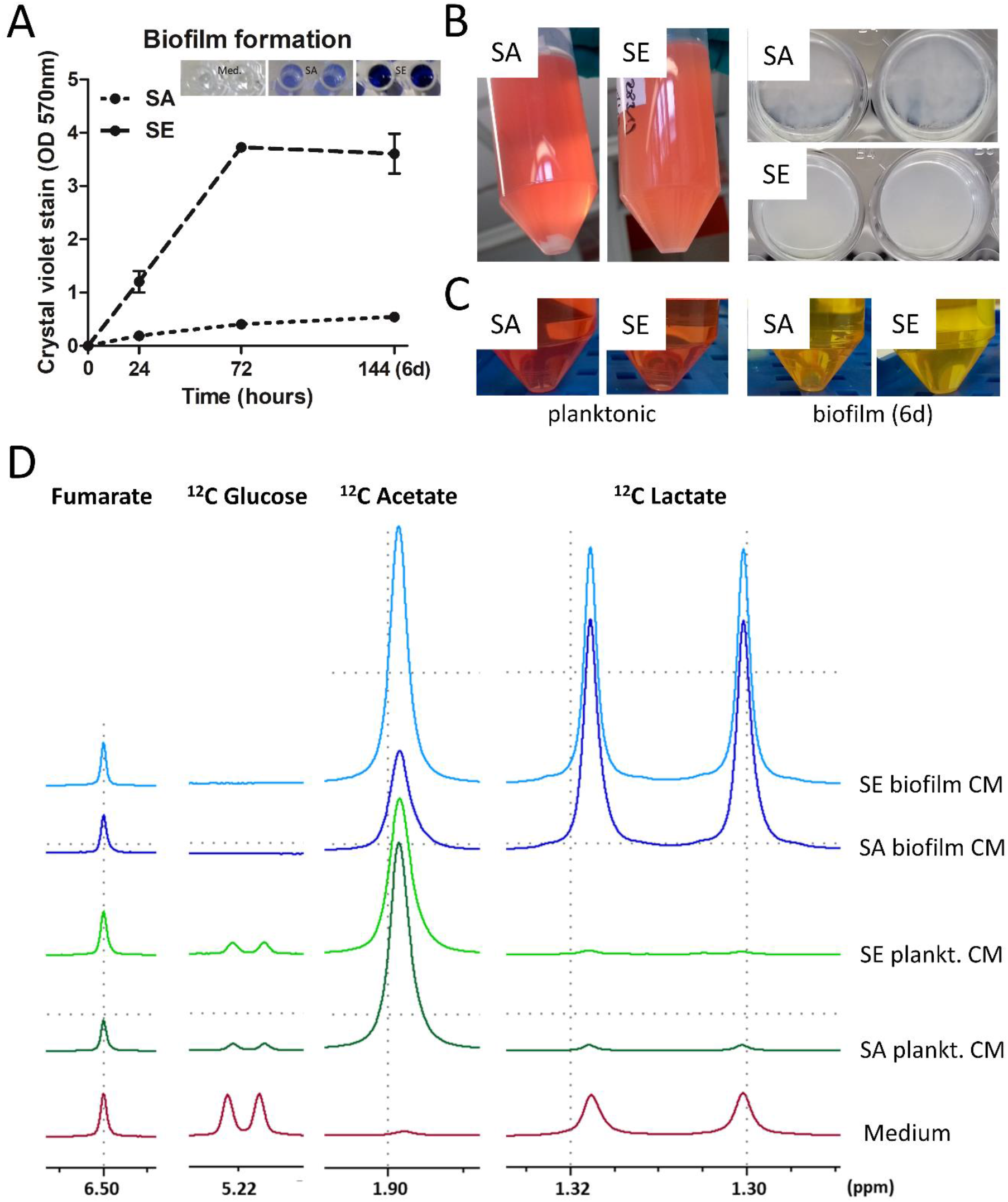
Biofilm formation of selected bacteria strains and preparation and characterization of conditioned media. *Staphylococcus aureus* (SA) strain ATCC 49230 (UAMS-1) and *Staphylococcus epidermidis* (SE) strain DSM 28319 (RP62A) were used in this study. Bacteria were freshly streaked out a day before experiment. To generate log-phase bacteria, 3-5 colonies were transferred in TSB, cultured for 3 hours and then adjusted to 10^6^ CFU per ml. A) *In vitro* biofilm formation of study strains. 100 μl of bacteria (10^5^ CFU/ well) in TSB with 1% glucose were transferred into 96 well plates and cultivated at 37°C for 24, 72 and 144 hours. TSB with 1% glucose was replaced every 24 hours. Biofilm formation was quantified by crystal violet staining (1% crystal violet in H2O). Data are presented as OD at 570 nm which is proportional to biofilm mass. n=2 experiments in triplicates, mean ± SD are shown. B) Pictures showing planktonic cultures after 24 hours of shaking and biofilm formation of static culture in 24 wells after 6 days. For generation of conditioned media (CM), bacteria (6*10^5^ CFU/ well/ ml) were cultured in growth media (DMEM high glucose + 10% FCS) without antibiotics at 37°C and 5% CO_2_. During biofilm formation media were replaced every 24 hours. C) Pictures of respective CM before pH adjustment. CM were harvested by centrifugation and sterile filtration after 24 hours of planktonic culture or the last 24 hours media of 6-days biofilm culture. D) ^1^H NMR spectra of CM were acquired using a 400 MHz Bruker spectrometer (Bruker Ultrashield^TM^ Plus 400). To allow comparison between different spectra, sodium fumarate (10 mM), dissolved in a 0.2 M phosphate buffer solution prepared with D2O (99.9%), was used as an internal standard. For comparison of different CM, spectra were adjusted to each other by equalizing the resonance of fumarate. Resonances of metabolites were then compared between the different CM. Shown are multiple display expansions of spectra of growth medium and CM for fumarate (singlet; δ = 6.50 ppm), ^12^C glucose (doublet; δ = 5.22 ppm), ^12^C acetate (singlet; δ = 1.90 ppm) and ^12^C lactate (doublet; δ = 1.31 ppm).

### All CM induce a pro-inflammatory macrophage immune response, which is less pronounced in biofilm CM

To evaluate potential differences in the activation of macrophages by either the planktonic or biofilm environments of SA and SE, we stimulated RAW 264.7 macrophages with the respective CM. As a control, we also stimulated cells with the TLR-2/-9 ligands Pam3CSK4 and CpG ODN (PC). Figure 2A shows that stimulation of macrophages with planktonic CM increased the surface protein expression of the immune activation markers TLR-2, MHC II and CD80 much stronger than the respective biofilm CM. This is in line with previous findings that the biofilm environment promotes immune cell tolerance. Protein levels of intracellular TLR-9, however, were only slightly increased with no major differences between planktonic and biofilm CM. Unexpectedly, gene expression analysis revealed that all CM induced the expression of the pro-inflammatory cytokine *Tnfa* to a similar extent. However, mRNA levels of the anti-inflammatory cytokine *Il10* were higher in biofilm CM. As TNF-α is associated with a pro-inflammatory (M1) and IL-10 with an anti-inflammatory (M2) macrophage phenotype, the ratio of the respective mRNA levels was used as an indicator of macrophage polarization. Here, the *Tnfa*/*Il10* ratio suggests an overall reduced pro-inflammatory macrophage polarization in biofilm compared to planktonic CM (Fig. 2B). Interestingly, in planktonic environments we observed an induction of *Ifnb* gene expression, which was highest for stimulation with SA planktonic CM (Fig. 2C). The corresponding protein analysis corroborated that TNF-α secretion was increased for all CM treatments, with the lowest concentration found for SA planktonic CM. Also, IL-10 release was increased on the protein level after CM treatment but in contrast to mRNA levels, no differences were observed between planktonic and biofilm environments (Supp. Fig. 1A). In line with the gene expression analysis, IFN-β protein levels were only increased after stimulation with planktonic CM (Supp. Fig. 1B). However, it has to be taken into account that high Protein A contents in SA planktonic CM might have interfered with the antibody-based detection and contributed to the cytokine concentrations measured in the supernatants of samples treated with SA planktonic CM.

**Figure 2.**
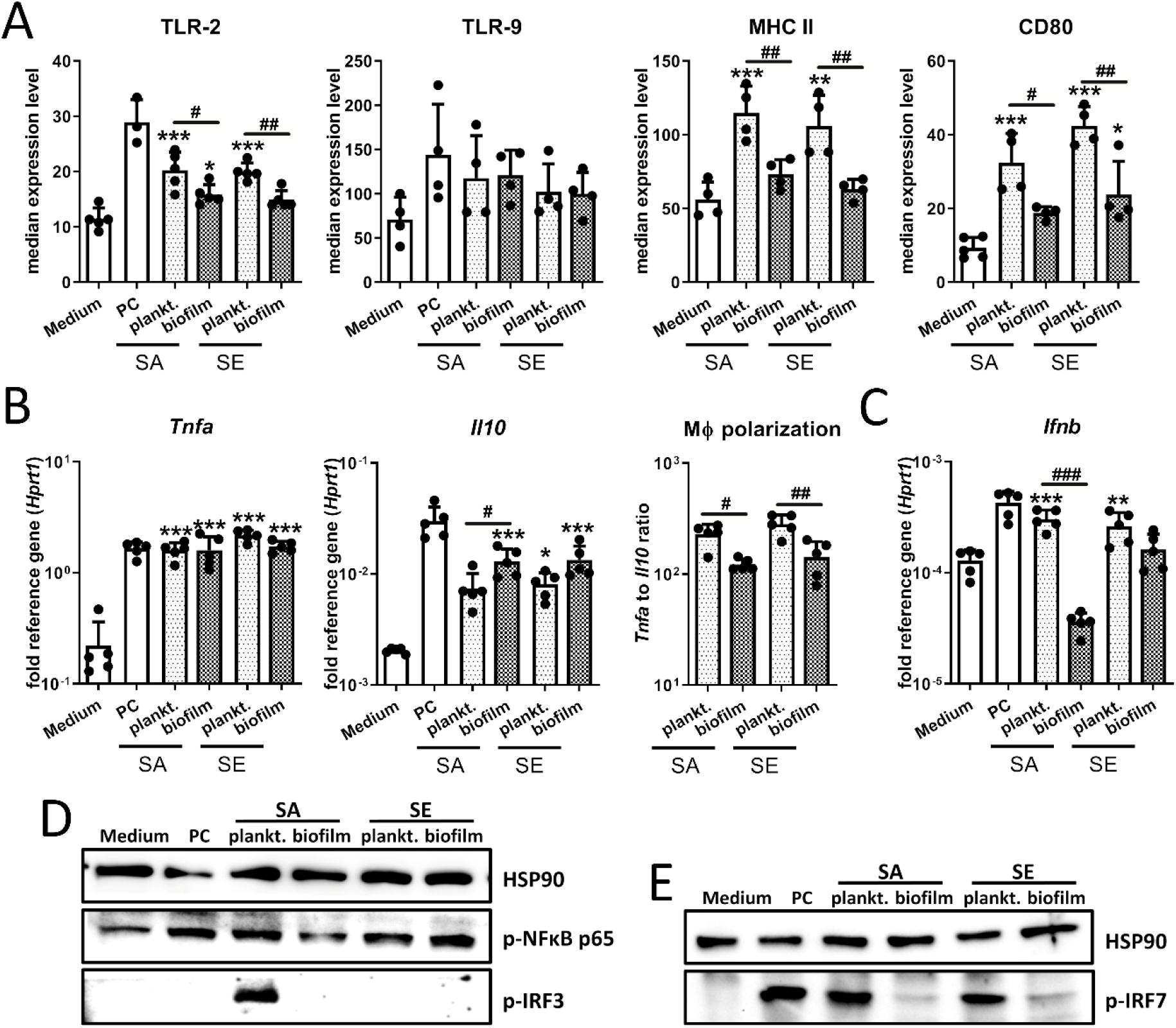
Macrophage immune activation after stimulation with conditioned media. RAW 264.7 cells were cultivated in CM 1:1 diluted in fresh growth media (DMEM high glucose + 10% FCS + 1% Pen/Strep) and immune response was investigated. A) Immune marker profile of macrophages. Cells were stimulated with CM for 20 hours and surface protein levels of TLR-2, MHC complex II (MHCII) and co-stimulatory protein CD80 as well as endosomal TLR-9 were measured by FACS analysis. Data are presented as median expression levels measured by fluorescence intensity. B+C) Gene expression analysis of relevant cytokines. Cells were stimulated with CM for 20 hours and mRNA levels of pro-inflammatory *Tnfa* and anti-inflammatory *Il10* (B) and *Ifnb* (C) were quantified by RT-qPCR. Ratio of *Tnfa* to *Il10* expression levels was used as indicator for macrophage polarization. Data are presented as relative gene expression of gene of interest related to the reference gene *Hprt1*. D+E) Activation of NF-κB, IRF3 and IRF7 signaling in macrophages. Cells were stimulated with CM for 4 hours and presence of phospho-NFκB p65 and phospho-IRF3 (D) or phospho-IRF7 (E) as activated forms of the transcription factors was visualized by Western Blot. HSP90 was used as loading control, n=3 experiments. For A-C: Data are presented as mean + SD and single values are shown as dots. n=4-5 experiments, p-values are calculated by Ordinary one-way ANOVA with post-hoc Bonferroni corrected multiple comparison. * is indicating significance against Medium, # is showing significance between respective planktonic and biofilm CM. * p<0.05, ** p<0.01, *** p<0.001; # p<0.05, ## p<0.01, ### p<0.001. PC: positive control (1 μg Pam3CSK4 + 100 nM CpG ODN).

On the level of signal transduction, we found an activation of the NF-κB pathway as indicated by increased levels of phosphorylated p65 (Fig. 2D), which is in line with the observed increase in *Tnfa* and *Il10* expression. While both SA and SE planktonic CM were able to trigger IRF7 phosphorylation, the master regulator of *Ifnb* induction, the IRF-3 pathway was only activated after treatment with SA planktonic CM. Also, the *Ifnb* gene expression caused by TLR activation was independent of phospho-IRF3 as only an activation of the IRF7 pathway was observed for PC (Fig. 2D + E). To rule out that the observed differences between planktonic and biofilm CM were due to cell viability, we stained for dead cells, but could not detect any differences between the treatment groups (Supp. Fig 1C). Thus, our data suggest that both planktonic as well as biofilm CM lead to a pro-inflammatory macrophage immune response, however, the expression of immune surface markers is less prominent after treatment with biofilm CM and the induction of an IFN-β response was limited to planktonic CM.

### Macrophage immune response is dominated by glycolytic metabolism with a shift towards increased mitochondrial activity in SE biofilm CM

As the metabolic state shapes the activity of immune cells and supports immune cell polarization, we analyzed the metabolic activity of RAW 264.7 macrophages upon CM treatment. In a first step we investigated the levels of L-lactate in the CM. As expected from the ^1^H-NMR results, only low levels of L-lactate were present in planktonic CM, whereas biofilm CM contained high amounts of bacterially derived L-lactate (Fig. 3A). Next, we measured L-lactate levels from RAW macrophage cultures after stimulation with the CM. Here, we detected increased L-lactate concentrations in the supernatants of all culture conditions. However, it has to be considered that the L-lactate generated by the bacteria during biofilm formation contributed to the L-lactate concentrations present in the supernatant of macrophages treated with biofilm CM (Fig. 3B). Thus, we corrected the measured L-lactate concentrations in the macrophage supernatants with the theoretical L-lactate concentration delivered by the respective CM 1:1 diluted in fresh cell culture media. The calculated L-lactate production by the macrophages indicated an increased aerobic glycolysis of the cells especially in the planktonic environment (Fig 3C). Aconitate decarboxylase 1 (ACOD-1, also known as IRG-1) catalyzes the reaction of cis-aconitate into the anti-inflammatory itaconate and thus interrupts the Krebs cycle, shifting cellular metabolic activity towards glycolysis [33, 34]. *Acod1* gene expression was induced by TLR-2/-9 ligands and CM-treated macrophages (Fig. 3D). A mitochondrial activity assay revealed that also mitochondrial activity was induced after stimulation with TLR-2/-9 ligands or CM. This was most pronounced in the biofilm environment of SE biofilm but comparatively low for treatment with TLR ligands (Fig. 3E). Total ATP levels remained similar across all treatment conditions (Fig. 3F).

**Figure 3.**
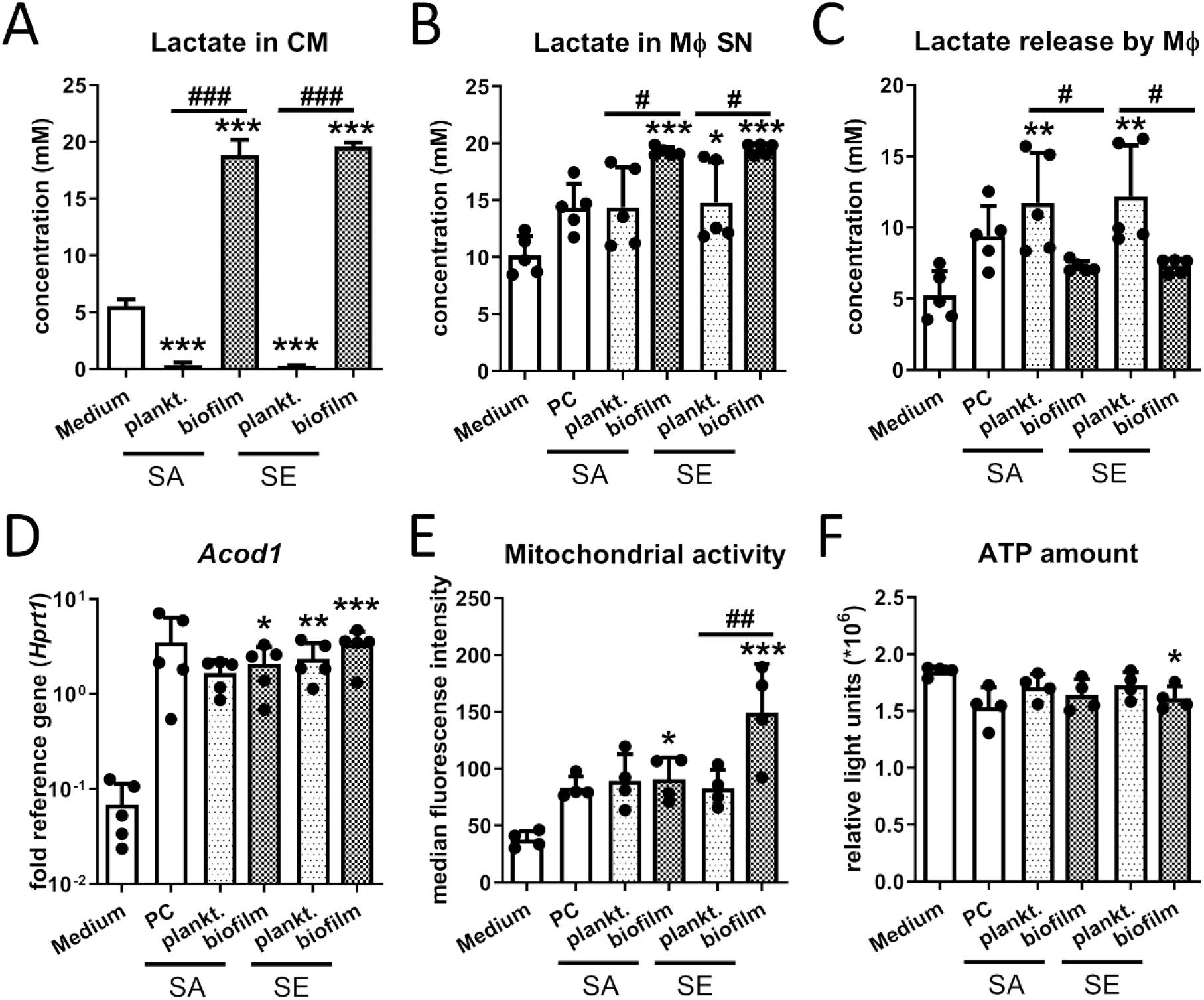
Metabolic changes in macrophages after stimulation with conditioned media. RAW 264.7 cells were cultivated in CM 1:1 diluted in fresh growth media (DMEM high glucose + 10% FCS + 1% Pen/Strep) and metabolic parameters were investigated. A) L-lactate amount in conditioned media. Bacteria were cultivated either in planktonic culture for 24 hours or biofilm culture for 6 days. CM were harvested 24 hours after last media exchange and L-lactate concentration was quantified by an enzyme-based assay. Data are presented as concentration (mM) measured by bioluminescent light release. B) L-lactate amount in supernatants of CM-treated macrophages. Cells were stimulated with CM 1:1 diluted in fresh growth media for 24 hours. Total L-lactate in the supernatant was quantified by an enzyme-based assay. Data are presented as concentration (mM) measured by bioluminescent light release. C) L-lactate release by macrophages after CM stimulation. Amount of L-lactate released by macrophages was calculated by total L-lactate concentration of macrophage supernatants minus the theoretical L-lactate concentration of CM 1:1 diluted in fresh growth medium. Data are presented as concentration (mM). D) Gene expression analysis of *Acod1*. Cells were stimulated with CM for 4 hours and mRNA levels of *Acod1* were quantified by RT-qPCR. Data are presented as relative gene expression of gene of interest related to the reference gene *Hprt1*. E) Mitochondrial activity of macrophages. Cells were stimulated with CM for 24 hours and mitochondrial activity was measured by FACS analysis using a membrane potential-dependent fluorescent dye. Data are presented as median mitochondrial potential measured by fluorescence intensity. F) ATP production by macrophages. Cells were stimulated with CM for 24 hours and total ATP content was measured in cell lysates by enzyme-based assay (CTG). Data are presented as relative light units measured by bioluminescent light release. For all: Data are presented as mean + SD and single values are shown as dots. n=4-5 experiments (for CTG assay mean of technical triplicates was included in statistics), p-values are calculated by Ordinary one-way ANOVA with post-hoc Bonferroni corrected multiple comparison. * is indicating significance against Medium, # is showing significance between respective planktonic and biofilm CM. * p<0.05, ** p<0.01, *** p<0.001; # p<0.05, ## p<0.01, ### p<0.001. PC: positive control (1 μg Pam3CSK4 + 100 nM CpG ODN).

Overall, our data corroborate that in a pro-inflammatory scenario macrophage metabolism mainly relies on aerobic glycolysis. Again, this was found to be more pronounced in the planktonic environment. Especially for SE, a shift towards increased mitochondrial activity become apparent in the biofilm environment, which generally is associated with an anti-inflammatory immune response.

### Low glucose and high lactate levels only show moderate but slightly different effects on macrophage cytokine response upon TLR-2/-9 stimulation

To investigate if the changes in the macrophage activity pattern were primarily caused by the observed shift in metabolite abundance in the biofilm environment, we stimulated RAW 264.7 macrophages with the TLR ligands Pam3CSK4 and CpG ODN (PC) in the presence of high (4.5 g/l) and low (1 g/l) glucose levels or different concentrations of lactate added to high glucose medium and investigated the resulting macrophage cytokine response and metabolic activity as described previously. Compared to eukaryotes where L-lactate is the predominant form, SA is able to produce both enantiomers, respectively [35]. Thus, we included D-lactate in our investigations. Figure 4A shows that there were only slight glucose concentration-dependent changes in the expression of *Tnfa* or *Il10*, which resulted in a statistically significant decrease in pro-inflammatory macrophage polarization (*Tnfα/Il10* ratio) in low glucose compared to a high glucose environment (Fig. 4A). The gene expression level of the glucose transporter *Slc2a1* was not affected by the amount of glucose in the medium (Fig. 4B). Different glucose concentrations also did not change basal or TLR induced mitochondrial activity (Fig. 4C). The increase in *Tnfa* mRNA levels observed upon TLR-2/-9 activation was further elevated by addition of extracellular L-lactate. However, this was not accompanied by a change in *Il10* induction. In sum, this led to a concentration-dependent increase of pro-inflammatory macrophage polarization by extracellular L-lactate upon TLR-2/-9 activation (Fig. 4D). Interestingly, increasing extracellular L-lactate levels further enhanced the mitochondrial activity observed upon stimulation with PC (Fig. 4E). Adding D-lactate to the medium had no effect on the gene expression levels of *Tnfa* and led to an only slightly but not significantly (p=0.1989 PC vs. PC+20mM) increase in induction of *Il10* mRNA levels not enough to change macrophage polarization (Fig. 4F). Eventually, the L-lactate present in the FCS of the cell culture medium has partially overshadowed the effects of D-lactate. This might equally explain, why the increasing D-lactate concentration led to an increase in mitochondrial activity upon TLR-2/-9 activation (Fig. 4G). We also investigated the effect of increasing extracellular acetate concentrations and found that gene expression levels of *Tnfa* only slightly (p=0.1046 PC vs. PC+20mM) but those from *Il10* significantly increased at high acetate concentrations, which did not result in an overall change in pro-inflammatory macrophage polarization (Supp. Fig. 2).

**Figure 4.**
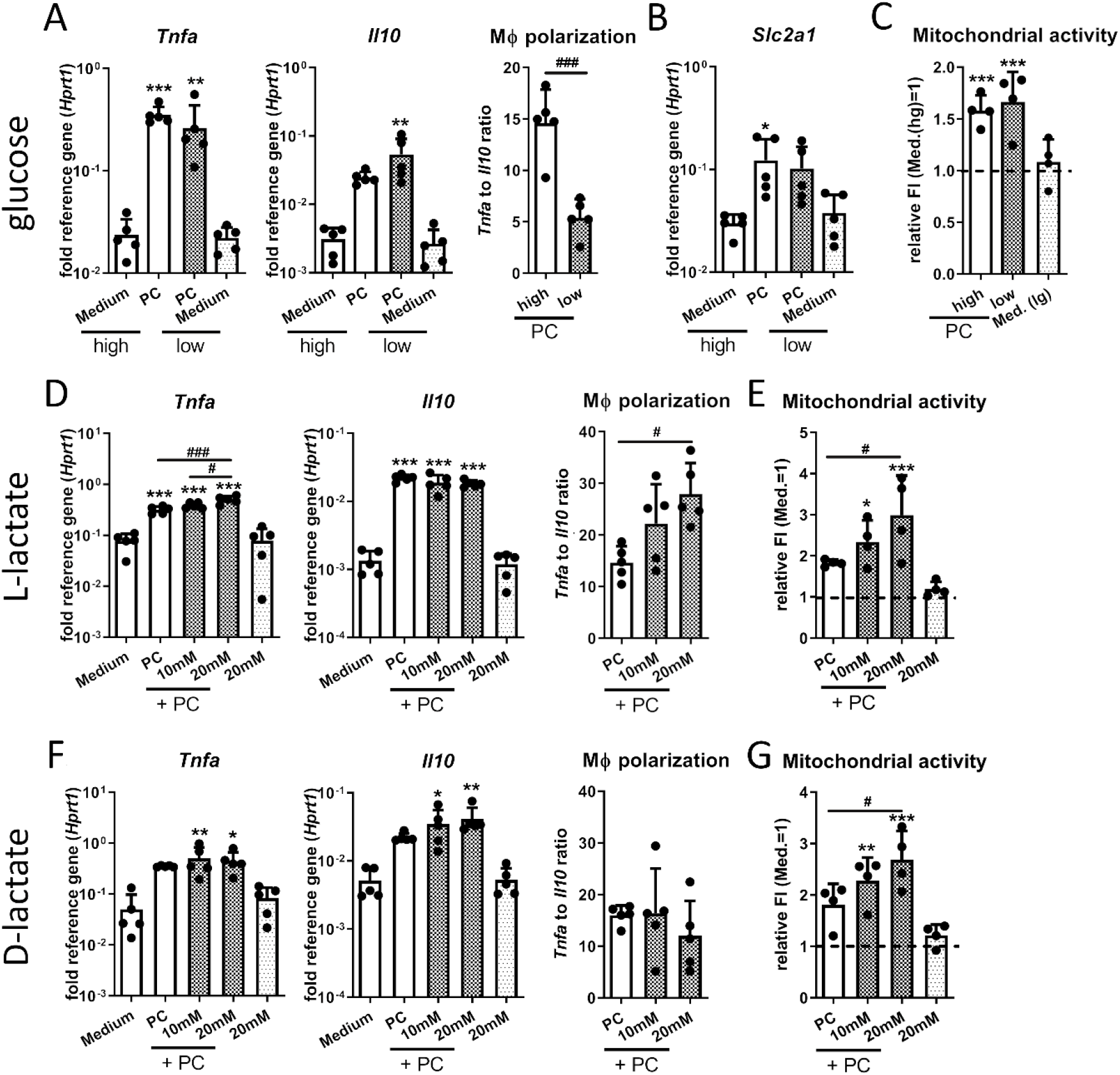
Effect of biofilm metabolite profile on macrophage TLR-2/-9 response. RAW 264.7 cells were stimulated with PC (positive control: 1 μg Pam3CSK4 + 100 nM CpG ODN) for 20 hours under low glucose or high lactate conditions and cytokine response and mitochondrial activity were evaluated. A-C) Effect of extracellular glucose concentration. Cells were stimulated by PC either in high (4.5 g/L) or low (1 g/L) glucose media. A+B) Gene expression analysis of pro-inflammatory *Tnfa* and anti-inflammatory *Il10* (A) and glucose transporter *Slc2a1* (B). Ratio of *Tnfa* to *Il10* expression levels was used as indicator for macrophage polarization. Data are presented as relative gene expression of gene of interest related to the reference gene *Hprt1*. C) Mitochondrial activity was measured by FACS analysis using a membrane potential-dependent fluorescent dye. Data are presented as normalized fluorescence intensity with medium sample set as 1. D+E) Effect of extracellular L-lactate concentration. Cells were stimulated by PC in media supplemented with different L-lactate concentrations (10 and 20 mM). D) Gene expression analysis of pro-inflammatory *Tnfa* and anti-inflammatory *Il10*. Ratio of *Tnfa* to *Il10* expression levels was used as indicator for macrophage polarization. Data are presented as relative gene expression of gene of interest related to the reference gene *Hprt1*. E) Mitochondrial activity was measured by FACS analysis using a membrane potential-dependent fluorescent dye. Data are presented as normalized fluorescence intensity with medium sample set as 1. F+G) Effect of extracellular D-lactate concentration. Cells were stimulated by PC in media supplemented with different D-lactate concentrations (10 and 20 mM). G) Gene expression analysis of pro-inflammatory *Tnfa* and anti-inflammatory *Il10*. Ratio of *Tnfa* to *Il10* expression levels was used as indicator for macrophage polarization. Data are presented as relative gene expression of gene of interest related to the reference gene *Hprt1*. H) Mitochondrial activity was measured by FACS analysis using a membrane potential-dependent fluorescent dye. Data are presented as normalized fluorescence intensity with medium sample set as 1. For all: Data are presented as mean + SD and single values are shown as dots. n=4-5 experiments, p-values are calculated by Ordinary one-way ANOVA with post-hoc Bonferroni corrected multiple comparison. * is indicating significance against Medium, # is showing significance between respective planktonic and biofilm CM. * p<0.05, ** p<0.01, *** p<0.001; # p<0.05, ## p<0.01, ### p<0.001.

In summary, our data indicate that low glucose or high lactate concentrations in the environment only have moderate effects on TLR-2/-9 mediated induction of *Tnfa* and *Il10* gene expression in macrophages. Although glucose and, partially, acetate can contribute to a more anti-inflammatory and L-lactate to a more pro-inflammatory macrophage immune response, the net effect remained comparatively small.

### Low glucose and high lactate concentrations have no effect on macrophage IRF3 mediated Ifnb gene expression induced by SA planktonic CM

We further investigated if different glucose and lactate concentrations affected the ability of macrophages to produce IFN-β in a SA planktonic environment. Therefore, we investigated IRF3 pathway activation and the subsequent induction of *Ifnb* gene expression to see if the low glucose and high lactate environment observed in SA biofilm CM could inhibit this pathway. We stimulated the cells with SA planktonic CM diluted in high or low glucose medium or added increasing concentrations of extracellular L- and D-lactate. Stimulation with SA biofilm CM was included for comparison. Figure 5A shows that glucose did not affect the increase in *Tnfa* and *Il10* mRNA levels observed for SA planktonic environment resulting in an unchanged macrophage polarization status (Fig. 5A). In addition, the induction of *Ifnb* gene expression and the IFN-β target gene *Isg15* remained unchanged by different extracellular glucose concentration (Fig. 5B). Consistently, activation of the NF-κB or IRF3 pathway by SA planktonic CM was independent of the extracellular glucose concentrations (Fig. 5C). Comparable results were observed for different L- (Fig. 5D-G) and D- (Fig. 5H-K) lactate concentrations. We could observe that the differences in induction of *Tnfa* and *Il10* gene expression and pro-inflammatory macrophage polarization between SA planktonic and SA biofilm CM were not triggered by the increased lactate concentrations characteristic for the biofilm environment (Fig. 5D+H). Furthermore, *Ifnb* mRNA and p-IRF3 protein levels upon stimulation with SA planktonic CM were not affected by addition of extracellular L- or D-lactate (Fig. 5E/F+I/J). The cells responded to the higher extracellular L- or D-lactate concentrations with a slightly increased induction of *Il6* mRNA levels upon stimulation with SA planktonic CM when compared with conditions without lactate addition (Fig 5 G+K). However, this increase did not reach the levels in the SA biofilm samples (Fig. 5G+K).

**Figure 5.**
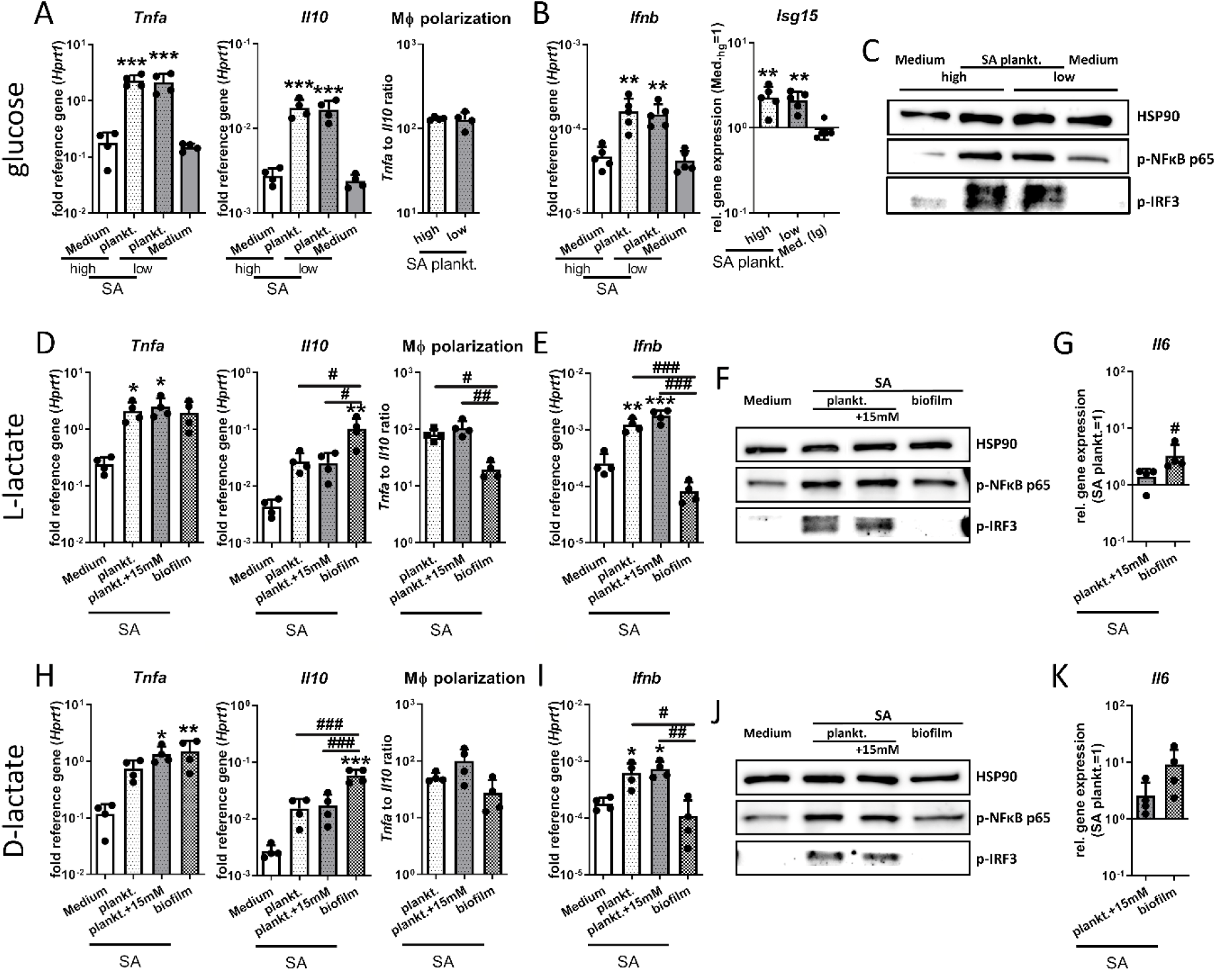
Effect of biofilm metabolite profile on IRF3 mediated *Ifnb* induction by SA planktonic CM. RAW 264.7 cells were cultivated in SA plankt. 1:1 diluted in fresh growth media (DMEM high glucose + 10% FCS + 1% Pen/Strep) with different extracellular glucose or lactate concentrations and immune response was investigated. A-C) Effect of extracellular glucose concentration. Cells were stimulated with SA plankt. CM 1:1 diluted either in high (4.5 g/L) or low (1 g/L) glucose media. A+B) Gene expression analysis of pro-inflammatory *Tnfa* and anti-inflammatory *Il10* (A) and *Ifnb* (B) after 20 hours of stimulation. Ratio of *Tnfa* to *Il10* expression levels was used as indicator for macrophage polarization. Data are presented as relative gene expression of gene of interest related to the reference gene *Hprt1*. C) Activation of NF-κB and IRF3 signaling in macrophages. Cells were stimulated for 4 hours and presence of phospho-NFκB p65 and phospho-IRF3 as activated forms of the transcription factors was visualized by Western Blot. HSP90 was used as loading control, n=3 experiments. D-G) Effect of extracellular L-lactate concentration. Cells were stimulated with SA plankt. CM 1:1 diluted in media ± 15 mM L-lactate or SA biofilm CM for comparison. D+E) Gene expression analysis of pro-inflammatory *Tnfa* and anti-inflammatory *Il10* (D) and *Ifnb* (E) after 20 hours of stimulation. Ratio of *Tnfa* to *Il10* expression levels was used as indicator for macrophage polarization. Data are presented as relative gene expression of gene of interest related to the reference gene *Hprt1*. F) Activation of NF-κB and IRF3 signaling in macrophages. Cells were stimulated for 4 hours and presence of phospho-NFκB p65 and phospho-IRF3 as activated forms of the transcription factors was visualized by Western Blot. HSP90 was used as loading control, n=3 experiments. G) Gene expression analysis of *Il6* after 20 hours of stimulation. Data are presented as relative gene expression normalized on SA plankt. CM. H-K) Effect of extracellular D-lactate concentration. Cells were stimulated with SA plankt. CM 1:1 diluted in media ± 15 mM D-lactate. G+H) Gene expression analysis of pro-inflammatory *Tnfa* and anti-inflammatory *Il10* (G) and *Ifnb* (H) after 20 hours of stimulation. Ratio of *Tnfa* to *Il10* expression levels was used as indicator for macrophage polarization. Data are presented as relative gene expression of gene of interest related to the reference gene *Hprt1*. J) Activation of NF-κB and IRF3 signaling in macrophages. Cells were stimulated for 4 hours and presence of phospho-NFκB p65 and phospho-IRF3 as activated forms of the transcription factors was visualized by Western Blot. HSP90 was used as loading control, n=3 experiments. K) Gene expression analysis of *Il6* after 20 hours of stimulation. Data are presented as relative gene expression normalized on SA plankt. CM. For A+B, D+E+G, H+I+K: Data are presented as mean + SD and single values are shown as dots. n=4-5 experiments, p-values are calculated by Ordinary one-way ANOVA with post-hoc Bonferroni corrected multiple comparison. * is indicating significance against Medium, # is showing significance between respective planktonic and biofilm CM. * p<0.05, ** p<0.01, *** p<0.001; # p<0.05, ## p<0.01, ### p<0.001.

Our data demonstrate that artificial low glucose and high lactate concentrations combined with the stimulation of macrophages with SA planktonic CM is not sufficient to induce a more anti-inflammatory macrophage immune response or to prevent IRF3 mediated *Ifnb* induction as seen in the respective biofilm environment.

## 4. Discussion

Biofilm formation is a major cause for chronic progression of implant-related bone infections. The biofilm environment is discussed to shift the immune reaction towards a more tolerogenic response that supports bacterial persistence. Macrophages play an important role in the early defense against invading bacteria and their pro-versus anti-inflammatory polarization is critical for effective bacterial clearance. In the present study we wanted to investigate if the biofilm metabolite environment characterized by low glucose and high lactate levels is a main factor determining the intensity and direction of the macrophage immune response in planktonic versus biofilm infectious situations. In addition, we included two relevant bacteria strains, *Staphylococcus aureus* and *epidermidis*. SA is highly virulent, produce a panel of toxins, is able to survive intracellularly and can form biofilms [12]. In contrast, the pathogenicity of SE mainly depends on biofilm formation [13]. To address this difference, we used the SE reference strain RP62A, which possesses a high *in vitro* biofilm formation capacity [32] and the SA strain UAMS-1, which originates from an osteomyelitis patient and shows only moderate biofilm formation on uncoated plastic surfaces [29, 31]. Independent of the different capacities in biofilm formation, the CM of both SA and SE shared similar characteristics with low glucose and high lactate levels in the biofilm CM, nicely representing glucose deprivation and lactate accumulation known for the local biofilm micromilieu [27, 28]. We cultivated RAW 264.7 macrophages in planktonic or biofilm CM generated from SA and SE cultures, respectively, and analyzed immune cell activation by measuring cell surface proteins, cytokine gene expression, metabolic activity and underlying signal transduction events. We showed that planktonic and biofilm environments both are able to elicit a predominantly pro-inflammatory immune response with increased glycolytic activity. However, this was less pronounced in biofilm CM and the increased gene expression levels of the anti-inflammatory cytokine IL-10 support this finding. Our data further indicate that only planktonic bacteria are able to initiate an IRF7 mediated IFN-β response, which was not detected in the respective biofilm environment. Interestingly, only in SA planktonic CM, *Ifnb* induction was also associated with IRF3 pathway activation. This can be explained by the profuse arsenal of virulence factors expressed by planktonic SA which causes more severe immune reactions than SE. Mimicking the metabolite profile of the biofilm environment with low glucose or high lactate concentrations had no effect on macrophage cytokine induction after TLR-2/-9 activation or stimulation with SA planktonic CM. In summary, our data indicate that the biofilm environment indeed elicits a less strong immune activation and supports a more anti-inflammatory macrophage phenotype compared to the respective planktonic environment. This was confirmed for SA as well as for SE. Further, our data clearly show that mimicking the biofilm metabolite environment during stimulation with relevant TLR ligands or planktonic CM is insufficient to shift the macrophage immune response towards the biofilm situation. Thus, our results suggest that ultimately, differentially released pathogenicity factors by the bacteria either growing planktonic or in biofilm are the central mediators that shape the resulting immune response.

In line with previous data from SA biofilm infection models [21, 22], biofilm CM of SA and SE induced a less pronounced upregulation of TLR-2 surface localization compared to planktonic CM. Despite the reduced surface levels of TLR-2, MHCII and CD80 in biofilm CM, planktonic as well as biofilm CM of both strains induced a comparable increase in NF-κB signaling that resulted in the expression of proportionate amounts of pro-inflammatory TNF-α. Equal or even increased TNF-α levels upon stimulation with supernatants generated from biofilm compared to planktonic SA cultures were also detected in human keratinocytes or fibroblasts [36, 37]. Our findings are in contrast to a similar study that found a suppression of pro-inflammatory macrophage activity by biofilm CM of SA which was mediated by KLF2 [38], a known negative regulator of NF-κB transcriptional activity [39]. The authors suggest that the observed increase in KLF2 expression might be caused by secretion of class II exotoxins such as α-hemolysin into the environment. However, the UAMS-1 strain used in our study is negative for α-hemolysin due to a mutation of the *hla* gene [40], which might explain the different results. Further, the authors focused on *Il6* and *Il1b* gene expression and did not check for *Tnfa* gene expression which might be differentially regulated. Our data suggest that the initial release of pro-inflammatory cytokine TNF-α from macrophages might be less important in the inefficient immune response against biofilms than the subsequent initiation of a T-cell response through MHCII mediated antigen presentation. As an impaired T cell response is discussed as one of the reasons behind chronicity of biofilm infections [7], effects of the biofilm environment on the macrophage/T-cell interaction should be investigated in more detail. Our experiments further demonstrated an induction of *Ifnb* gene expression in macrophages upon stimulation with planktonic CM of SA and SE. For SA infections, induction of an IFN-β response has been described previously [41]. Here, we showed that also SE is able to trigger IFN-β production. The direct comparison of *Ifnb* induction by planktonic and biofilm environments revealed that, this specifically happens in the presence of a planktonic environment and is not triggered by the corresponding biofilm environment. Planktonic CM strongly induced TLR signaling that could have been one cause for the activation of the IRF7 pathway detected in these environments. For SA planktonic CM an additional activation of IRF3 was observed, which can be a downstream target of cytosolic nucleic acid sensor pathways such as the cGAS-STING pathway [42]. A recent study showed that indeed STING-IRF3 signaling is involved in IFN-β induction upon infection of macrophages with live SA [43]. Another study showed that also SA biofilms can lead to a STING-dependent IFN-β induction in macrophages via the release of c-di-AMP due to bacterial lysis [44]. This is in contrast to our findings as we did not detect IRF3 activation and *Ifnb* induction in the biofilm environment which could be caused by the use of different SA strains, varying culturing conditions and experimental setups. In virus infections, IFN-β interferes with cellular proliferation and induces the production of ISGs that impairs viral replication [45]. It is becoming increasingly clearer that IFN-β has important but controversial functions in bacterial infections [46, 47]. The fact that *Ifnb* was dominantly expressed after incubation with planktonic bacteria that usually can be cleared through the immune system suggests that a focus should be set on investigating its role in the course of chronic implant-related bone infections. Metabolic analysis revealed that especially in the planktonic environments, the macrophage immune response is dominated by aerobic glycolysis. However, it could be observed that the cells started to enhance their mitochondrial activity upon stimulation with CM, which was most pronounced for CM derived from SE biofilm cultures. Increased OxPhos activity is associated with a more anti-inflammatory M2 macrophage polarization [26] and might play a role in chronic biofilm infections [28]. Our finding for SE biofilm CM is in line with a recent study, where a shift towards OxPhos activity was shown in monocytes over the time course of an orthopedic biofilm infection. Furthermore, the authors showed that inhibiting OxPhos *in vivo* by a nanoparticle-based delivery of oligomycin restored an effective pro-inflammatory monocyte immune response and reduced biofilm burden [48]. Extracellular lactate was found to be associated with an inhibitory effect on the pro-inflammatory immune response of macrophages [49–51]. In a recent study, the group of Tammy Kielian compared the effects of biofilm derived L- and D-lactate on the production of anti-inflammatory IL-10 using mutant SA strains deficient in L- and D-lactate production [35]. Their data suggest that biofilm-derived lactate is responsible for an increased IL-10 synthesis by myeloid-derived suppressor cells (MDSCs) and macrophages via inhibition of histone deacetylase 11 (HDAC11) and an unchecked *Il10* promotor activity. Although we detected high amounts of bacterial lactate only for biofilm CM, in our experimental setup, a strong *Il10* induction could be detected for both, planktonic and biofilm CM. In the biofilm CM, induction of *Il10* gene expression was higher than in the respective planktonic environment which might be due to its higher lactate concentration. As we did not detect bacterial lactate in planktonic CM, we suggest that the mechanisms behind *Il10* induction may differ between planktonic and biofilm environments. In addition, independently of bacterial derived lactate concentrations, planktonic CM was able to induce *Il10* gene expression as a consequence of increased TLR signaling.

We further evaluated the effect of extracellular low glucose or high lactate levels on macrophage polarization upon TLR-2/-9 stimulation. Most studies have investigated the impact of extracellular lactate as a product of aerobic glycolysis triggered by LPS mediated TLR-4 signaling [52]. In these studies, high lactate concentrations were associated with a suppression of a pro-inflammatory macrophage responses [49, 51], which was also seen for TLR-2 stimulation by Pam3Cys [50]. Conversely, we detected an increase of *Tnfa* mRNA levels with increasing extracellular L-lactate concentrations resulting in a more pro-inflammatory macrophage polarization upon TLR-2/-9 activation. In comparison to our setup, the other studies used either higher lactate concentrations (up to 100 mM) or lactate pre-incubation before LPS treatment which might have led to different results. Further, it is possible that the effects of lactate on the macrophage immune response vary between different TLR ligands. In addition, we observed that addition of extracellular lactate dose-dependently enhanced mitochondrial activity after stimulation with TLR-2/-9 ligands, which after a longer time period of several days might lead to a metabolically induced switch towards a more anti-inflammatory response. Compared to lactate, the impact of extracellular glucose levels on a TLR mediated immune response has been investigated to a lesser extent. High extracellular glucose levels were linked to LPS-induced inflammasome activation, pyroptosis and IL-1β production in macrophages [53, 54]. In our setting, we only observed minor effects of extracellular glucose levels on *Tnfa* and *Il10* gene expression levels after TLR-2/-9 stimulation. Nonetheless, the ratio between the two indicated that macrophage polarization shifted towards a less pro- and more anti-inflammatory response at low glucose levels. Mitochondrial activity, however, remained unaffected. In a paper investigating the effects of different glucose concentrations on LPS-mediated immune responses of macrophages from non-diabetic and diabetic mice [55], it was found that the effect of extracellular glucose concentrations on the LPS response was getting stronger over time. After 24 hours, the authors detected only slight changes in cytokine release of healthy macrophages, whereas higher glucose concentrations decreased TNF-α cytokine levels after 7 days of LPS stimulation. Furthermore, we investigated the effects of different extracellular glucose and lactate concentrations on the macrophage cytokine response upon stimulation with SA planktonic CM. Low glucose or addition of lactate had no effect on induction of *Tnfa* gene expression in response to SA planktonic CM nor increased *Il10* mRNA levels like in the SA biofilm CM. This clearly differed from our results observed for our bacterial CM which were shown by ^1^H-NMR to contain low glucose and high lactate concentrations under biofilm conditions. Our data suggest that mimicking biofilm metabolite conditions in a planktonic environment by reducing glucose or adding lactate was not sufficient to shift the macrophage polarization towards the biofilm situation. Therefore, the difference in the macrophage activation profile is primarily dependent on further substances from the CM and not on the metabolite levels. Low glucose or high lactate levels also did not affect IRF3 activation and the subsequent induction of *Ifnb* gene expression upon stimulation with SA planktonic CM. This again indicates that the metabolite profile of the biofilm environment does not prevent the IRF3 mediated *Ifnb* induction. It rather seems that IFN-β production requires an additional bacterial stimulus that is present under planktonic conditions but missing in the biofilm environment. This is not unexpected, as the induction of bacterial genes associated with biofilm formation and metabolic adaption is often accompanied by a downregulation of virulence factors predominantly expressed in the planktonic lifestyle [56, 57]. Identifying these immunogenic, bacterial mediators that discriminate between planktonic and biofilm environments will help to develop future treatment options to strengthen the immune response in chronic implant-related bone infections.

## Conflict of Interest

The authors have no relevant financial or non-financial interests to disclose.

## Author contribution

ES was responsible for study conception and design, acquisition, analysis and interpretation of data and wrote the manuscript. TE and FVK participated in data acquisition, analysis and interpretation. MSC was involved in data interpretation and critically revised the manuscript. KFK supervised the study, contributed to data interpretation, helped to draft the manuscript and critically revised the manuscript. All authors read and approved the final manuscript.

## Funding

Elisabeth Seebach was funded by the Physician Scientist Program of the Medical Faculty of Heidelberg University.

## Acknowledgements

We would like to thank Katharina Draxel and Stella Cavicchioli for their help with the experiments. Furthermore, we thank Gabriele Sonnenmoser for technical assistance.

## Additional information

Supplementary material accompanies the manuscript, which is available in a separate file. This manuscript is available online in the pre-print version https://doi.org/10.1101/2021.07.26.453923.

## Notes

### Competing Interest Statement

The authors have declared no competing interest.

### Summary of Updates

Previous manuscript version was divided into two. This manuscript version focuses on the macrophage immune response data. Data analysis and statistical evaluation were revised and additional experiments are included.

