## Supplementary material for "Bacterial and metabolic factors of staphylococcal planktonic and biofilm environments differentially regulate macrophage immune activation": Suppl. material

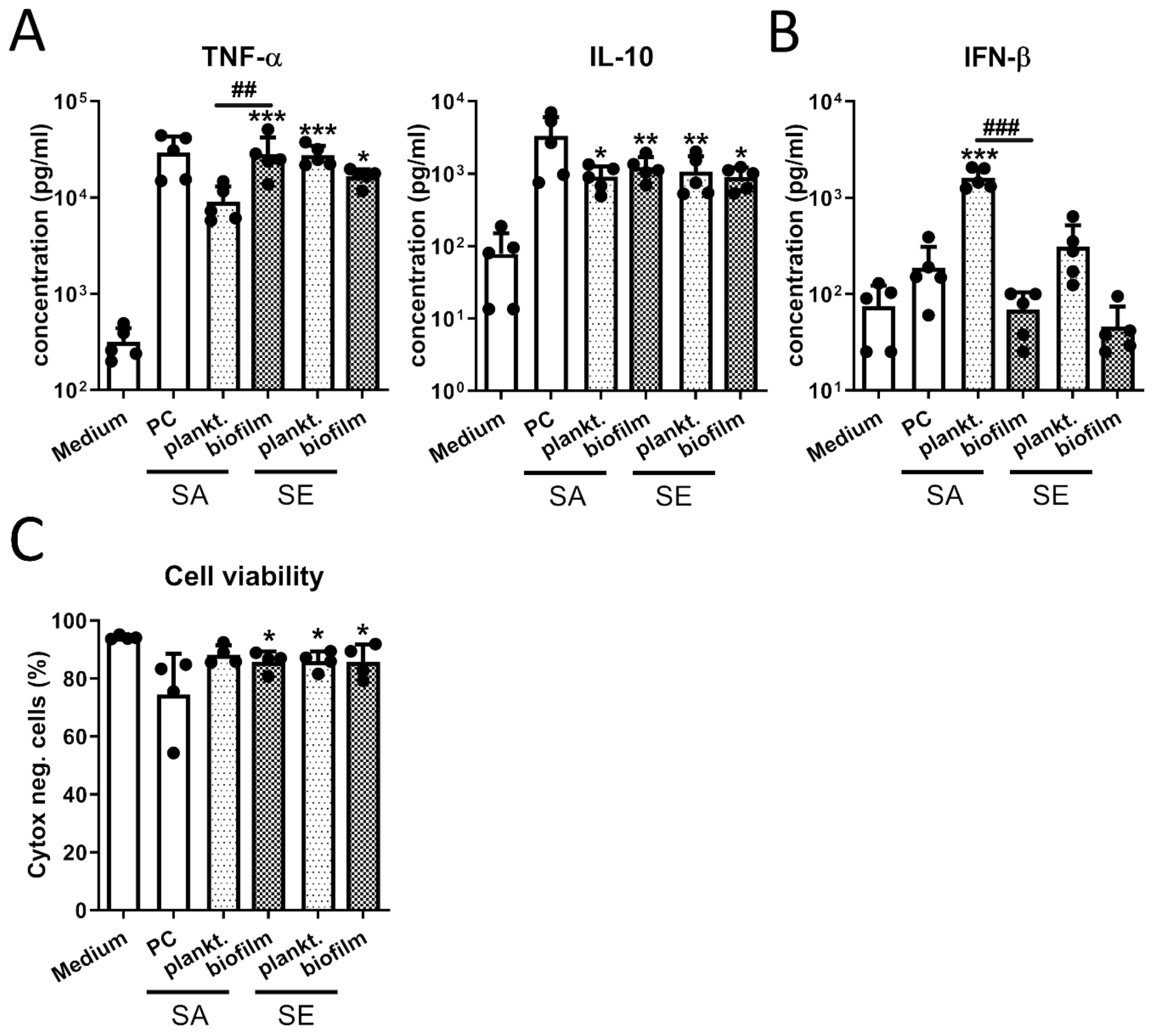


Suppl. Figure 1. **Protein concentrations of cytokines released by macrophages and cell viability upon stimulation with CM.** RAW 264.7 cells were cultivated in CM 1:1 diluted in fresh growth media (DMEM high glucose + 10% FCS + 1% Pen/Strep) and cytokine release was analyzed. Cells were stimulated with CM for 20 hours and concentrations of pro-inflammatory TNF-α and anti-inflammatory IL-10 (A) as well as IFN-β (B) were quantified in the supernatant by cytometric bead array (CBA; LegendPlex). Data are presented as absolute concentration (pg/ml). C) Viability of macrophages after CM stimulation. Cells were stimulated with CM for 20 hours and cell viability was measured by Cytox-staining and FACS analysis. Percentage of Cytox-negative (living) cells are shown. For all: Mean + SD are shown with single values as dots. n=4-5 experiments, p-values are calculated by Ordinary one-way ANOVA with post-hoc Bonferroni corrected multiple comparison. * is indicating significance against Medium, # is showing significance between respective planktonic and biofilm CM. * $p<0.05$, ** $p<0.01$, *** $p<0.001$; # $p<0.05$, ## $p<0.01$, ### $p<0.001$. PC: positive control (1 µg Pam3CSK4 + 100 nM CpG ODN).

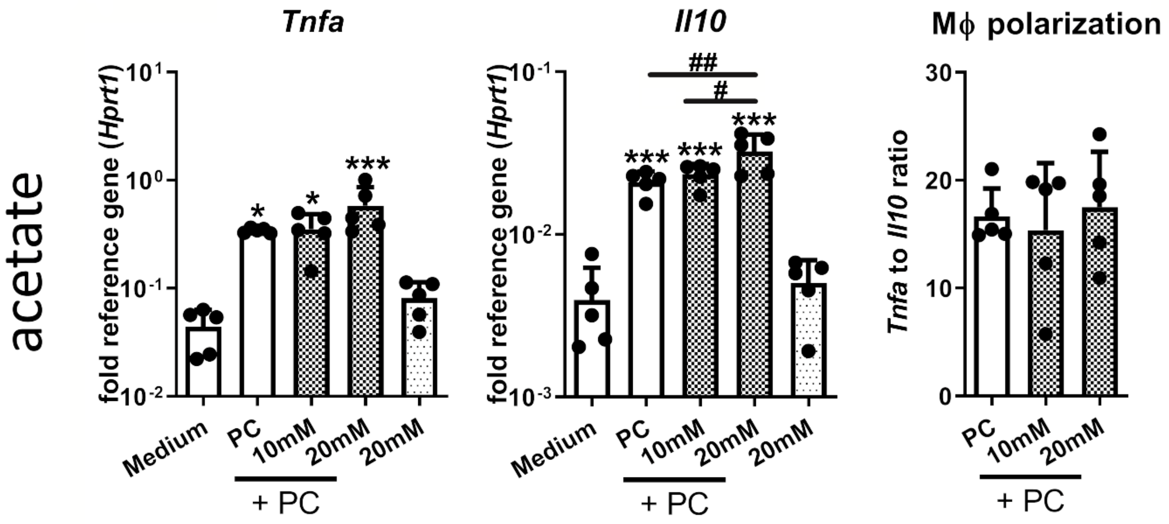


Suppl. Figure 2. **Effect of bacterial derived acetate on macrophage TLR-2/-9 response.** RAW 264.7 cells were stimulated with PC (positive control: 1 µg Pam3CSK4 + 100 nM CpG ODN) for 20 hours with different extracellular acetate concentrations (10 and 20 mM) added to the medium and immune response was evaluated. Gene expression analysis of pro-inflammatory *Tnfa* and anti-inflammatory *Il10*. Ratio of *Tnfa* to *Il10* expression levels was used as indicator for macrophage polarization. Data are presented as relative gene expression of gene of interest related to the reference gene *Hprt1*. Mean + SD are shown with single values as dots. n=5 experiments, p-values are calculated by Ordinary one-way ANOVA with post-hoc Bonferroni corrected multiple comparison. * is indicating significance against Medium, # is showing significance between respective planktonic and biofilm CM. * p<0.05, ** p<0.01, *** p<0.001; # p<0.05, ## p<0.01, ### p<0.001.
